# Generalized Method of Moments improves parameter estimation in biochemical signaling models of time-stamped single-cell snapshot data

**DOI:** 10.1101/2022.03.17.484491

**Authors:** Wu John, William CL Stewart, Jayaprakash Ciriyam, Das Jayajit

## Abstract

**Motivation:** Ordinary differential equations are commonly used to model the sub-cellular dynamics of average values of proteins and mRNAs. New single-cell technologies provide cell-to-cell differences in protein/mRNA abundances that allow for the evaluation of higher order moments. However, using this additional information to improve parameter estimation can be challenging since the magnitudes of single-cell abundances can vary widely between proteins/mRNA.

**Results:** We employ Generalized Method of Moments (GMM) and Particle Swarm Optimization to address the above challenges in mechanistic modeling of signaling kinetics data. Using synthetic data from linear and non-linear models, we show that the proposed method improves parameter estimation and enables construction of approximate confidence intervals. Furthermore, our approach exploits parallel computation to scale with increasing data size and dimensions. We apply our software CyGMM to estimate parameters in a linear ODE model for publicly available longitudinal single-cell cytometry data for CD8+ T cells. Our results demonstrate substantial improvements for modeling data from single-cell cytometry and RNA-seq experiments.

**Availability:** We also make freely available our estimation software CyGMM written in C++ on github (https://github.com/jhnwu3/CyGMM).

**Supplementary information:** Supplementary data are available at *Bioinformatics* online.

## 1 Introduction

Ordinary differential equations (ODEs) are commonly used to model the sub-cellular dynamics of proteins and mRNA (Rohrs et al. (2019); Loskot et al. (2019); La Manno et al. (2018)). Usually, ODEs describe the deterministic dynamics of average concentrations, and estimation of model parameters from experimental data is an important step towards building biological models. The task of estimating parameters is challenging due to a variety of reasons (Stewart (2018); Ashyraliyev et al. (2009); Raue et al. (2009)), one of which is that the number of model parameters is larger than that of the available data. Recent developments of experimental techniques for the longitudinal measurements of transcripts and proteins at the single cell level (e.g., single-cell RNA seq(Saliba et al. (2014)), CyTOF for proteins(Mukherjee et al. (2017); Spitzer and Nolan (2016)) appear to alleviate this problem. For any time t, the cell-to-cell variability in the copy numbers of proteins and RNA arises from two sources: variation present at the pre-stimulus state (*aka* extrinsic noise), and variation that arises from the stochasticity of biochemical reactions (*aka* intrinsic noise) (Swain et al. (2002); Das and Jayaprakash (2018)). Especially, in developing models to describe signaling kinetics when the protein abundances are large, extrinsic noise is known to play a significant role (Feinerman et al. (2008)) and single cell protein signaling kinetics can be well approximated by ODEs. The single-cell data provide information about the probability distribution of different proteins due to extrinsic noise across cells. This leads to the availability of more observables, e.g., the various moments of the protein numbers, than the number of parameters in an ODE model. However, single cell abundances of different protein species can vary by orders of magnitudes (Hui et al. (2017); Milo and Phillips (2015)) thus mean values and higher moments of these abundances can differ by many orders of magnitudes. Hence, the challenge here is to estimate the parameters reliably in this case given that the observed values can vary over many orders of magnitude. In this paper, we address this problem by devising a method based on the Generalized Method of Moments (GMM) to models of interacting proteins in which extrinsic noise dominates. GMM, a method used widely in econometrics (Hansen (1982); Hall (2004)), has been used previously to estimated parameters in simple models describing transcription and translation with intrinsic noise but no extrinsic noise fluctuations. We use GMM in conjunction with the Particle Swarm Optimization technique (Poli et al. (2007)) to minimize the cost function that depends on the difference between the sample and model moments. We discuss some of the potential challenges and show that our procedure can deal with these challenges both for linear and nonlinear ODEs, and for published CyTOF data. We also make available software (CyGMM) to implement the method.

## 2 System & Methods

We use a flexible ODE model to describe single cell signaling kinetics of protein abundances 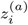, where *i* labels each of the *n* proteins in a single cell indexed by *a*. The abundances evolve in time through *r* biochemical signaling reactions according to

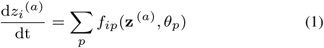

Here, *f*_*ip*_(**z** ^(*a*)^, *θ*_*p*_) is the propensity of the *p*^th^ reaction affecting the abundance of the *i*^th^ protein. Furthermore, **z** ^(*a*)^ is an *n*-dimensional vector of protein abundances, and *θ*_*p*_ is the rate constant for the *p*^th^ biochemical signaling reaction. Note that the ODE model in (1) allows for both linear and nonlinear dynamics (See Algorithm Implementation for further details).

The rest of this section is organized as follows. First we describe our parameter estimation approach (i.e. GMM) using first moments. Second, we show how our approach can be used to incorporate higher order moments. Third, we discuss the construction of approximate 95% CI’s for individual rate constants.

### 2.1 First Order Moments & Weight Matrices

Because cytometry experiments do not track individual cells over time, we observe instead the protein abundances of clonal cell populations at specific moments in time. We refer to these time-stamped observations of the dynamic process as *snapshot* data, and the analysis of such data permits estimation of the rate constants of an ODE model. To compare the statistical properties of competing estimators, we first generate synthetic snapshot data with ***θ*** known (we refer to this as the *ground truth*) according to the ODE model in (1). Then, using the same ODE model, we analyze the synthetic data assuming ***θ*** is unknown. For simplicity of exposition, we first describe our approach for estimators of that rely on differences in mean protein abundances. This simple setting is often useful and pragmatic, especially when the number of biochemical signaling reactions is less than the number of distinct proteins measured (i.e. *r < n*). Later, we will describe a systematic way to combine higher order moments.

For a single cell indexed by *a*, let **y** ^(*a*)^ denote the protein abundances generated with known rate constants at time *t*. Similarly, for a single cell indexed by *b*, let **x**(***θ***) ^(*b*)^ denote the protein abundances generated with hypothesized rate constants ***θ*** at time *t*. Recall that the initial cell-to-cell differences in protein abundances arise from extrinsic noise, which we model with a multivariate log-normal distribution (see Supplementary Information for details).

As is customary with GMM, we begin by constructing a set of *so called* moment conditions, where the difference in moments is zero when ***θ*** equals the true rate constants (denoted ***θ***^(*g*)^). In this case, those conditions are based on differences in mean protein abundances, and the conditions are given by

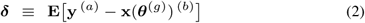

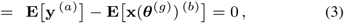

where the expectations in (3) are taken with respect to the multivariate log-normal distribution used to model the extrinsic noise in both clonal cell populations. From snapshot data across *N* cells, we can use the difference in sample means to estimate (3)

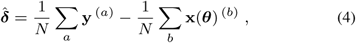

where we assume here (without loss of generality) that both clonal cell populations contain *N* cells. Recall that, for large samples, 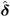 will be close to 0 whenever ***θ*** is close to the true rate constants. Hence, the GMM estimate of ***θ*** is defined implicitly as

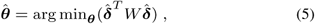

where *W* is a symmetric, positive definite weight matrix. The positive definite condition ensures that the distance in (5) (*aka* the cost function) is well-defined. When *W* is the identity matrix, the GMM estimator of ***θ*** is the standard least squares estimator (denoted 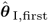).

Furthermore, Hansen (Hansen (1982)) showed that when the true rate constants are known, the best weight matrix is any matrix proportional to

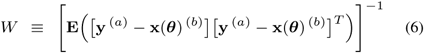

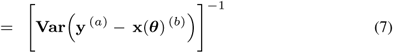

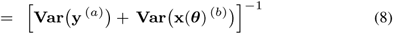

Note that Eq. (7) follows from Eq. (6) because ***δ*** = 0 when the true rate constants are known, and Eq. (8) follows from Eq. (7) because the clonal cell populations are independent (i.e. the covariance between **y** ^(*a*)^ and **x**(***θ***) ^(*b*)^ is zero). For 1 ≤ *i, j* ≤ *n*, we can compute the *ij*^th^ element of **Var**[**y**^(*a*)^] as,

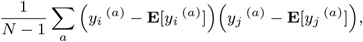

The weight matrix given by (8) is best because the corresponding GMM estimator (denoted 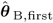) has the smallest variance among all GMM estimators of ***θ*** (cite). Here, **E**[*y*_*i*_ ^(*a*)^] is evaluated using 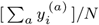. Analogous equations are used to compute **Var**[**x**(***θ***^(*g*)^)^(*b*)^] and **E**[*x*_*i*_(***θ***^(*g*)^) ^(*b*)^].

Because *W* in (8) depends on ***θ***, an iterative approach is often used. For instance, one can start by estimating ***θ*** without weights (i.e. setting *W* = *I* and using standard least squares). Then, one can apply Eq. (8) to the snapshot data at time *t* to estimate unequal weights. Of course, this new estimate of *W* can then be used with equation (5) to arrive at a second estimate of ***θ***. And so the iterative process continues until some preset number of iterations is reached, or until a convergence criterion is met. Interestingly, one can avoid the iterative process by using a nonparametric estimate of *W* (Lück and Wolf (2016))

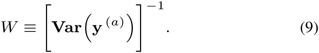

Note that, because we are generating synthetic snapshot data from a given ODE model, the weight matrix in (9) is proportional to, and slightly less efficient than, the best weight matrix in (8).

In practice however, matrix inversion in (8) and (9) may not always work (see Supplementary Information for more details). If this happens, then we can usually still improve upon the standard least squares estimator by setting the off-diagonal elements in the variance-covariance matrices of (8) or (9) to zero. This in turn yields a diagonal weight matrix with diagonal elements denoted *w*_*i*_, and the corresponding GMM estimator (dentoed 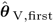) can be expressed as a weighted sum of the squared differences in mean abundances

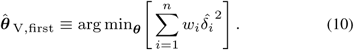

### 2.2 Higher Order Moments & Multiple Time Points

So far, we have described our GMM approach as it applies to first order moments, but our approach generalizes to the inclusion of higher order moments as well. To include second order moments (or variances), consider the following moment conditions:

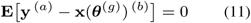

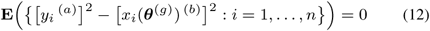

where the corresponding estimates are denoted 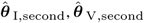 and 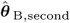, respectively. The expectation values for the clonal populations a and b are computed by assuming independence of protein abundances between the populations. Furthermore, the inclusion of *mixed* moments (or covariances) follows from the cross-product analog of (12). The corresponding estimates are denoted 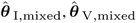 and 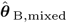, respectively.

Finally, our GMM approach also generalizes to the analysis of snapshot data with observations across multiple time points. For ease of exposition let’s revisit equation (10), which is only concerned with means, and let’s consider snapshot data with observations at time points *t*_1_ and *t*_2_. To estimate ***θ*** in this setting, we must find

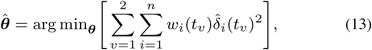

where (as before) *w*_*i*_ and *δ*_*i*_ both depend on the time *t*_*v*_ when the *snapshot* are taken.

### 2.3 Constructing Confidence Intervals

An attractive feature of our software (CyGMM) is its ability to construct approximate 95% confidence intervals (CI’s) for individual rate constants (denoted *θ*_*p*_). By using a resampling approach to construct CI’s (which we describe below), we avoid computationally expensive gradient approximations that often arise when CI’s are based on maximum likelihood. Specifically, we use the standard interval (nonparametric) bootstrap procedure (Tibshirani and Efron (1993)), and the asymptotic theory of GMM estimators, to construct approximate 95% CI’s. Hansen (Hansen (1982)) showed that for large samples, GMM estimators are approximately normally distributed with variance Σ, and this variance is easily estimated from a large number of bootstrap resamples. Therefore, we resampled the available snapshot data 50 times with replacement, computed 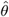 for each resample, and used these values to compute 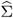. Then, we estimate the standard deviation of 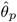 (denoted 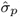) using the square root of the diagonal elements of 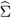. Finally, to construct approximate 95% CI’s, we set the lower bound of the interval to 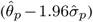 and the upper bound to 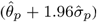. Note that, because *θ* ∈ {0, 1}^*r*^ where *r* is the number of biochemical signalling reactions, any CI that is not contained in [0,1] was truncated at 0 or 1, accordingly. Note that coverage, which is the probability that a CI contains *θ*_*p*_, was estimated from a total of 5,000 snapshot data sets, 100 synthetic data sets (i.e. replicates) with 50 bootstrap resamples per replicate.

## 3 Algorithm Implementation

We used a set of coupled ODE’s, both linear and nonlinear, to simulate and analyze snapshot data. For the linear ODE’s, the time-evolution of protein abundances required matrix exponentiation (Mukherjee et al. (2017)); whereas for the nonlinear ODE’s, we use the Runge-Kutta Cash-Karp nonlinear solver (see https://www.boost.org/) to evolve protein abundances over time. We then estimated parameters in an ODE model that describes published (Krishnaswamy et al. (2014)) (available at https://dpeerlab.github.io/dpeerlab-website/dremi-data.html) longitudinal single-cell CyTOF dataset for mouse naive CD8+ T cells stimulated by CD3/CD28 antibodies.

### 3.1 ODE Linear Model for Simulation

As a first demonstration of the method, we consider a set of protein species *X*_1_,…, *X*_*q*_ (*q* = 3, 6, or 10) where the species interact with each other via first order biochemical reactions in a single cell. We assume all species are measured at the same time *t*. The reaction networks for *q* = 3, *q* = 6, and *q* = 10 are shown schematically in Figures 1A, 1B, and 1C, respectively.

**Figure 1.**
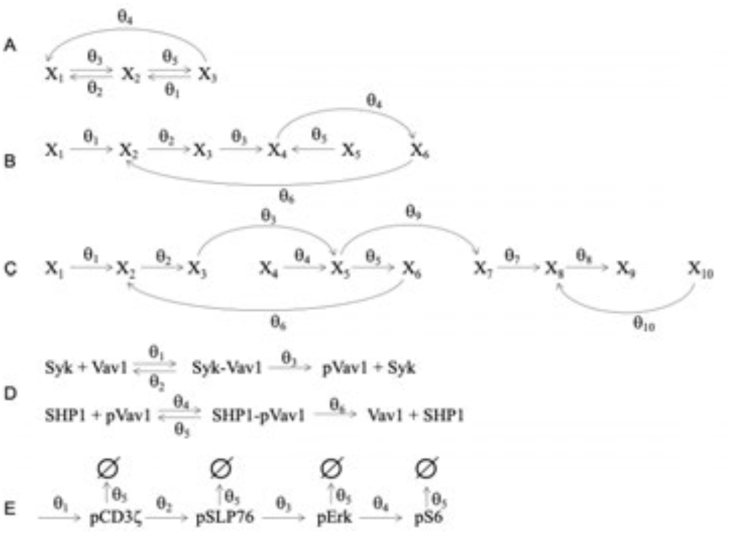
Schematic diagram of linear and nonlinear reaction models. Diagrams A-D are used to generate synthetic snapshot data. Diagram E was used to analyze published single-cell CyTOF data.

The protein abundances 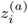 follow the linear ODEs given by

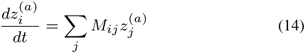

where, the elements of the *M* matrix are given by zero or linear sums of the reaction rates *θ*_*p*_. We assume no net creation or destruction of protein species, therefore, ∑_*i*_ M_*ij*_ = 0. The solution of Eq. (14) is given by,

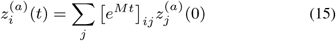

The above solution is used to evaluate means and higher moments for the data generated from the linear ODEs. The matrix exponentiation in (15) is carried out numerically using routines available in R and C++ (see available code at: gethub link for details). The rate constants *θ* are chosen between 0 and 1 (Table 1), and the times considered are 0 and 1.5, as determined by the range of true rate constants.

**Table 1.** Synthetic Snapshot Data Simulation Details For each model we generated synthetic snapshot data for 5,000 cells. Rate constants (as shown) are rounded to the nearest percent. The number of proteins for each model (from top to bottom) are: 3, 6, 10, and 6, respectively.

| Model | $\theta_1$ | $\theta_2$ | $\theta_3$ | $\theta_4$ | $\theta_5$ | $\theta_6$ | $\theta_7$ | $\theta_8$ | $\theta_9$ | $\theta_{10}$ |
| --- | --- | --- | --- | --- | --- | --- | --- | --- | --- | --- |
| Linear | 0.28 | 0.84 | 0.44 | 0.04 | 0.30 |  |  |  |  |  |
| Linear | 0.52 | 0.06 | 0.10 | 0.90 | 0.06 | 0.07 |  |  |  |  |
| Linear | 0.52 | 0.06 | 0.10 | 0.90 | 0.06 | 0.07 | 0.18 | 0.75 | 0.34 | 0.23 |
| Nonlinear | 0.10 | 0.10 | 0.95 | 0.17 | 0.05 | 0.18 |  |  |  |  |

### 3.2 ODE Nonlinear Model for Simulation

We consider a simplified model of biochemical reactions that describe early time signaling kinetics in mouse Natural Killer (NK) cells stimulated by ligands cognate to activating CD16 and inhibitory Ly49H receptors. We model the kinase Syk bound to activating receptor-ligand complex and the phosphotase SHP-1 bound to inhibitory receptor-ligand complex (Fig. 1D). The enzyme Syk phosphorylate a key signaling protein Vav1 where phosphorylated Vav1 (pVav1) induce activation of NK cells. The enzyme SHP-1 dephosphorylate pVav1 and inhibit NK cell activation. In our model, Syk and SHP1 react with the substrate protein Vav1 and phosphorylated Vav1 (pVav1) following the reactions below. Activating and inhibitory NK cell receptors are not included explicitly in this model, instead, Syk and SHP-1 represent Syk and SHP-1 proteins bound the the activating and inhibitory receptors, respectively. The reactions in the model are similar to the zero-order ultrasensitivity model proposed by Goldbeter and Khoshland (Goldbeter and Koshland (1981)) which is widely used to describe various biochemical systems. The abundances for the proteins Syk, Vav1, Syk-Vav1, SHP1, SHP1-pVav1 and pVav1 evolve in time with a set of nonlinear ODEs describing mass-action kinetics for the above reactions. We assume all the six above species are measured at any time t. The ODEs are shown in Supplementary Information, and the parameter values for the ground truth model (see Table 1) are taken from the parameters used in (Das (2010)). The ODEs are solved numerically using a Runge-Kutta Cash-Karp nonlinear solver in C++ at selected time points: 0s, 0.5s, 2s, 10s, 20s, and 30s to generate the synthetic data.

### 3.3 ODE Model for CyTOF Data Describing T-cell Signaling Kinetics

We employed a linear ODE model (Fig. 1E) to describe T cell signaling kinetics involving the protein species pCD3*ζ*, pSLP-76, pErk, and pS6 assayed by time stamped snapshot CyTOF data (Krishnaswamy et al. (2014)). The ODE’s minimally describe phosphorylation of tyrosine residues in CD3*ζ* chains due to crosslinking of CD3 and CD28 co-receptors in mouse naive CD8+ T cells by their cognate antibodies. Production of the phosphorylated form of CD3*ζ* or pCD3*ζ* leads to phosphorylation of SLP76 subsequently induces phosphorylation of Erk. Phosphorylated form of Erk (or pErk) produces phosphorylation of the protein S6. Dephosphorylation of the phosphorylated forms of the proteins by phosphatases is captured by the first order decay of these proteins with the rate *θ*_5_. The model considered here is a substantially reduced version of the processes that generate the above signaling events (Krishnaswamy et al. (2014)). The ODEs are shown in the Supplementary Information. The CyTOF experiments measured protein abundances for pCD3*ζ*, pSLP76, pERK, and pS6 in 1500 mouse naive CD8+ T cells stimulated by CD3/CD28 antibodies at time points t=0, 0.5min, 1 min, 2 min, 4min and 8 min. We estimated the rate parameters *θ*_1_, *θ*_2_, *θ*_3_, *θ*_4_, and *θ*_5_ for time interval *t*_1_ = 1min to *t*_2_ = 2min. We find that the values of the estimated rate parameters depend on the time interval considered indicating time dependence of the rates in T-cell signaling kinetics. The means and higher order moments for the above protein abundances were calculated by removing negative abundances and a small fraction of cells (approx. 6%) with very large abundances (i.e. observed values that were in excess of three standard deviations from the mean). The ODEs in the model cannot be solved by (15) and instead were solved numerically using the Runge-Kutta Cash-Karp solver in C++ as described in the previous subsection.

## 4 Results

As a proof of principle, we first show results from the linear ODE model with a single time point involving either 3, 6, or 10 proteins. In this setting, across a wide range of weight matrices, we show that CyGMM provides consistent estimates of *θ*. We use mean squared error (MSE) to compare competing estimators, because MSE is minimized when an estimator is both accurate (i.e. has low bias) and precise (i.e. has low variance). Specifically, the MSE of 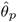 is defined as its variance 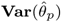 plus the square of its bias 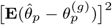, for *p* = 1,…, 5. For each *θ*_*p*_, we also compute an approximate 95% CI and its coverage (i.e., the probability that the CI covers the true rate constant *θ*_*p*_). Note that, when one computes a CI, there is no guarantee that the interval will contain the true rate constant. In fact, the best one can ever hope for is that 19 out of 20 times, the 95% CI covers the truth. Therefore, to confirm the validity of our approximate CI’s, we computed their coverage probabilities from 100 replicates. After presenting results for the linear ODE model, we show results for the biologically inspired (and more complex) nonlinear ODE model. In the nonlinear setting snapshot data across multiple time points are generated and analyzed. Here (as with the linear ODE model), we use MSE to compare estimators that use different weight matrices and different sets of moment conditions.

### 4.1 Estimation with Linear ODE’s

MSE along with variance, squared bias, coverage, and the average CI lengths are shown for the linear ODE model with 3 proteins and 5 rate constants (Table 2). In terms of MSE, the estimator that combines first, second and mixed moments (i.e. means, variances, and covariances) with optimal weights (denoted 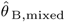), performs the best. The estimator that uses the sub-optimal weights as defined in Eq. (10), denoted 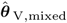, is a close second. When one uses equal weights, the squared bias is large and the coverage is poor (see Table 2 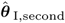 and 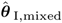). However, the estimation does improve when covariances are incorporated (i.e. 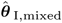 had lower MSE and better coverage than 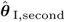). Therefore, while the inclusion of higher order moments can improve the estimation, it is more important to incorporate optimal (or at least sub-optimal) weights.

**Table 2.** MSE and Coverage (Linear Model with 3 Proteins) MSE, variance, and squared bias are shown (rounded to 4 decimal places) for three different GMM estimators. The first two estimators: 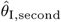, and 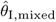 employ equal weights, but different numbers of moment conditions. The third estimator 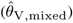 uses all moment conditions and weights that vary inversely in proportion to the relevant variances [see Eq. (10)]. The fourth estimator 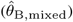 uses all moment conditions with optimal weights. Also shown, are the average values of individual rate constants across 100 replicates, the coverage probabilities for approximate 95% CI’s, and the average lengths of the corresponding intervals. Note that 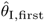 is not shown because some rate constants were unidentifiable.

| Estimator | Rate | Avg | MSE | Var | Bias<sup>2</sup> | Coverage | Length |
| --- | --- | --- | --- | --- | --- | --- | --- |
| $\hat{\theta}_{I,\text{second}}$ | $\theta_1$ | 0.173 | 0.0185 | 0.0100 | 0.0085 | 0.85 | 0.346 |
| | $\theta_2$ | 0.802 | 0.1127 | 0.0296 | 0.0831 | 0.54 | 0.613 |
| | $\theta_3$ | 0.533 | 0.0601 | 0.0588 | 0.0013 | 0.91 | 0.707 |
| | $\theta_4$ | 0.146 | 0.0487 | 0.0239 | 0.0248 | 0.88 | 0.448 |
| | $\theta_5$ | 0.307 | 0.1119 | 0.0947 | 0.0172 | 0.81 | 0.709 |
| $\hat{\theta}_{I,\text{mixed}}$ | $\theta_1$ | 0.253 | 0.0030 | 0.0025 | 0.0005 | 0.84 | 0.161 |
| | $\theta_2$ | 0.668 | 0.0436 | 0.0162 | 0.0274 | 0.69 | 0.432 |
| | $\theta_3$ | 0.340 | 0.0269 | 0.0162 | 0.0107 | 0.79 | 0.410 |
| | $\theta_4$ | 0.070 | 0.0018 | 0.0010 | 0.0008 | 0.84 | 0.107 |
| | $\theta_5$ | 0.312 | 0.0073 | 0.0072 | 0.0001 | 0.89 | 0.258 |
| $\hat{\theta}_{V,\text{mixed}}$ | $\theta_1$ | 0.274 | 0.0028 | 0.0028 | 0 | 0.96 | 0.204 |
| | $\theta_2$ | 0.809 | 0.0174 | 0.0166 | 0.0008 | 0.94 | 0.436 |
| | $\theta_3$ | 0.420 | 0.0073 | 0.0068 | 0.0005 | 0.95 | 0.315 |
| | $\theta_4$ | 0.042 | 0.0006 | 0.0006 | 0 | 0.99 | 0.106 |
| | $\theta_5$ | 0.300 | 0.0051 | 0.0051 | 0 | 0.94 | 0.260 |
| $\hat{\theta}_{B,\text{mixed}}$ | $\theta_1$ | 0.279 | 0.0014 | 0.0014 | 0 | 0.92 | 0.040 |
| | $\theta_2$ | 0.844 | 0.0003 | 0.0003 | 0 | 0.95 | 0.049 |
| | $\theta_3$ | 0.443 | 0.0001 | 0.0001 | 0 | 0.95 | 0.039 |
| | $\theta_4$ | 0.038 | 0.0001 | 0.0001 | 0 | 0.94 | 0.031 |
| | $\theta_5$ | 0.299 | 0.0027 | 0.0027 | 0 | 0.90 | 0.092 |

Next, we applied CyGMM to a more complicated linear ODE model with 6 proteins, and 6 rate constants. A similar pattern manifests (Table 3), in that 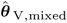 outperforms 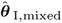, and they both outperform 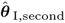. As before, incorporating higher order moments and unequal weights leads to improved estimation. Note that we also considered a linear ODE model involving 10 proteins and 10 rate constants. CyGMM estimated all 10 parameters well (see Supplementary Information), using all moment conditions and near-optimal weights [see Eq. (10)]. Therefore, given the linear ODE models explored here, CyGMM provides accurate (i.e. low bias) and precise (i.e. low variance) estimation, especially when higher order moments are used in conjunction with optimal (or near-optimal) weights. Although exact solutions for the nonlinear ODE’s are not possible, we wanted to see which (if any) of the main findings from the linear setting transferred to the nonlinear setting.

**Table 3.** MSE (Linear Model with 6 Proteins) The MSE, variance, and squared bias (rounded to 4 decimal places) are shown for three different GMM estimators. The first two estimators: 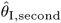, and 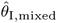 employ equal weights, but different numbers of moment conditions. The third estimator 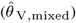 uses unequal weights to combine all moment conditions (see Eq. 10 for more details). Also shown are the average values of individual rate constants across 100 replicates, Because we could not compute the best weight matrix in this setting, 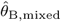 is not shown.

| Estimator | Rate | Avg | MSE | Var | Bias<sup>2</sup> |
| --- | --- | --- | --- | --- | --- |
| $\hat{\theta}_{I, \text{second}}$ | $\theta_1$ | 0.700 | 0.0460 | 0.0120 | 0.0340 |
| | $\theta_2$ | 0.113 | 0.0087 | 0.0060 | 0.0027 |
| | $\theta_3$ | 0.135 | 0.0072 | 0.0062 | 0.0010 |
| | $\theta_4$ | 0.787 | 0.0195 | 0.0074 | 0.0121 |
| | $\theta_5$ | 0.079 | 0.0059 | 0.0053 | 0.0006 |
| | $\theta_6$ | 0.091 | 0.0061 | 0.0058 | 0.0003 |
| $\hat{\theta}_{I, \text{mixed}}$ | $\theta_1$ | 0.539 | 0.0268 | 0.0263 | 0.0005 |
| | $\theta_2$ | 0.069 | 0.0005 | 0.0004 | 0.0001 |
| | $\theta_3$ | 0.116 | 0.0016 | 0.0014 | 0.0002 |
| | $\theta_4$ | 0.752 | 0.0419 | 0.0210 | 0.0209 |
| | $\theta_5$ | 0.053 | 0.0004 | 0.0004 | 0 |
| | $\theta_6$ | 0.078 | 0.0017 | 0.0017 | 0 |
| $\hat{\theta}_{V, \text{mixed}}$ | $\theta_1$ | 0.518 | 0.0004 | 0.0004 | 0 |
| | $\theta_2$ | 0.064 | 0.0006 | 0.0006 | 0 |
| | $\theta_3$ | 0.105 | 0.0004 | 0.0004 | 0 |
| | $\theta_4$ | 0.899 | 0.0004 | 0.0004 | 0 |
| | $\theta_5$ | 0.058 | 0.0005 | 0.0005 | 0 |
| | $\theta_6$ | 0.074 | 0.0004 | 0.0004 | 0 |

### 4.2 Estimation with Nonlinear ODE’s

In contrast to the three linear ODE models, our nonlinear model is computationally much more demanding and is based on an existing biochemical signaling network. Not surprisingly, it brings new statistical challenges as well. For example, after generating synthetic snapshot data for larger, and larger samples sizes with our nonlinear model, we found that our best estimate of *θ*_2_ converges to zero, which means that consistent estimation of the second rate constant is not possible since *θ*_2_ = 0.10. In general, when there are estimation problems on the boundary, researchers may want to consider estimating those parameters from some additional source of information. Note that even when the value of *θ*_2_ is incorrectly specified (e.g. *θ*_2_ = 0.05 and *θ*_2_ = 0.25) the estimates for the remaining rate constants are still quite good (see Table 6 Supplemental Information for more details). Furthermore, in the nonlinear scenario, we see from (Figure 2) that the estimation is improved when the analysis includes variances and covariances. The most notable reduction in MSE (mean squared error) occurrs in 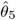.

**Figure 2.**
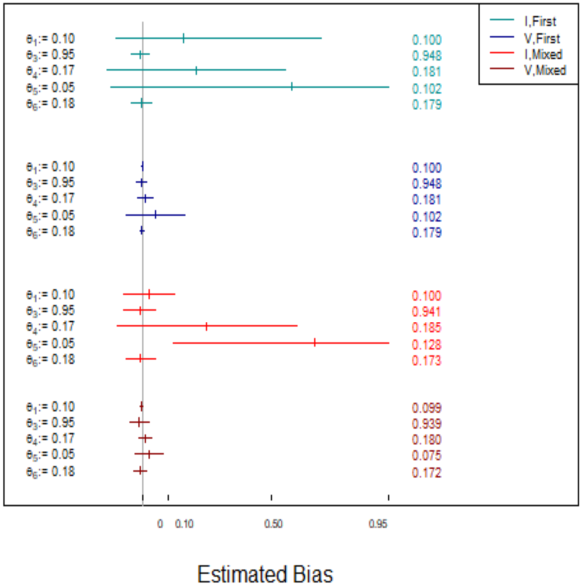
Near optimal weights improve estimation, and higher moments can reduce bias. Four GMM estimators of the nonlinear model are shown. Two estimators consider differences in means (light blue and dark blue), and two other estimators consider differences in all first, second, and mixed moments (light red and dark red). Lighter colors indicate the use of equal weights; darker colors indicate the use of unequal (i.e. near optimal) weights. The simulation truth is shown in the left margin, and for each rate constant, the average estimate across replicates is in the right margin.

**Figure 3.**
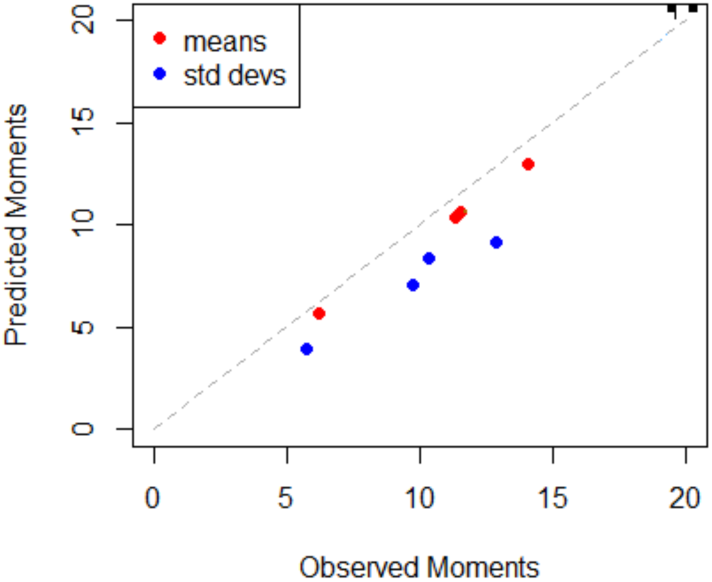
Model predictions agree with published CyTOF data. Scatter plot of predicted means versus observed means at 2 minutes taken from single-cell CyTOF dataset (red), and predicted standard deviations versus observed standard deviations at 2 minutes (blue).

### 4.3 Parameter Estimation Using Single Cell CyTOF Dataset for T-cell Signaling Kinetics

We used our software package, CyGMM to analyze the published CyTOF data of pCD3*ζ*, pSLP76, pERK, and pS6. The 5 estimated rate constants for the CyTOF data for *t*_1_=1 min and *t*_2_=2min and their corresponding CI’s are shown in Table (4). The relatively large widths of the CI’s for *θ*_1_ and *θ*_4_ indicate high variability, which is consistent with the corresponding pairwise contour plots (see Figure 4, Supplemental Information). The predicted values of means and variances at *t*_2_=2min show excellent to reasonable agreement with their experimental counterparts. As seen in Figure (3), the predicted means are close to the observed means (i.e. within 9%), whereas the predicted variances (i.e. standard deviations) are all further away from the observed variances.

**Figure 4.**
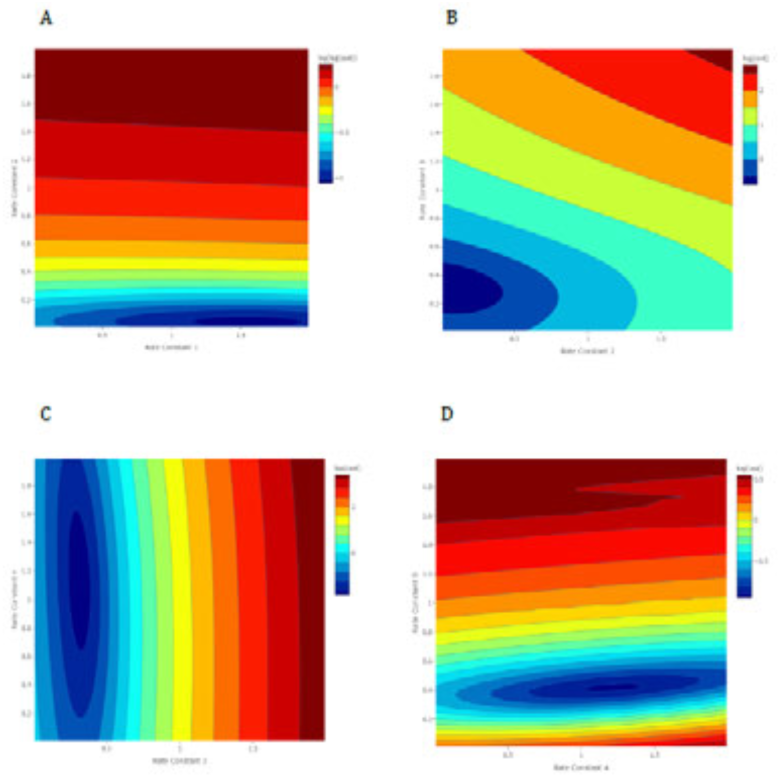
Pairwise contour plots are consistent with a unimodal high dimensional cost function. The 5-dimensional cost function fit to published CyTOF data considered differences in means and variances and used near optimal weights. For consecutive pairs of rate constants evaluation points ranged over a 2-dimensional grid with the remaining parameters fixed at their estimated values.

**Table 4.** Parameter Estimates with 95% CI’s Point estimates for each rate constant, along with its corresponding approximate 95% CI are shown. CI limits on the boundary of the parameter space are indicated with an asterisks (*).

| Rate Constant | Estimate | Lower Limit | Upper Limit |
| --- | --- | --- | --- |
| $\theta_1$ | 1.54 | 0.97 | 2* |
| $\theta_2$ | 0.04 | 0* | 0.13 |
| $\theta_3$ | 0.31 | 0.27 | 0.36 |
| $\theta_4$ | 1.12 | 0.90 | 1.35 |
| $\theta_5$ | 0.38 | 0.33 | 0.44 |

## 5 Conclusion

We developed a software package CyGMM to estimate rate parameters using time-stamped high dimensional single-cell cytometry (CyTOF) snapshot data in ODE models describing deterministic signaling kinetics with extrinsic noise. CyGMM applies GMM, a method developed and widely used in econometrics, to systematically combine means and higher moments of protein abundances whose magnitudes can span several orders of magnitudes and thus addresses the challenge of estimating greater number of model parameters than available data. Our approach leverages parallel computation for simulating single cell signaling kinetics, optimization of multidimensional cost functions via PSO, and computation of CIs for the estimated parameters. This represents a substantial improvement over existing approaches where GMM has been applied for smaller gene regulatory models devoid of extrinsic noise fluctuations or parameters in signaling kinetic models are estimated using bulk data. Application of CyGMM to synthetic data generated from linear and non-linear ODEs show its ability to estimate parameters and CIs accurately.

A challenging task in GMM is to compute the optimal weight factors to combine moments especially in large dimensions. We used optimal and non-optimal but computationally efficient weight factors and showed that our non-optimal weight factors provide almost as precise estimations as the optimal weights for the signaling models (see for example Table 2). This is probably because most of the weight tends to go to the differences in means, and because the correlations between protein abundances are usually not that large.

### Limitations and future directions

Estimating highly codependent parameters whose values can vary dramatically without significantly changing the signaling kinetics can be a challenge for CyGMM (especially when the ODE’s are nonlinear). For example, in the non-linear model we studied here, *θ*_1_, *θ*_2_, and *θ*_5_ represents such a parameter group. While identifying such groups may not be easy, a working solution could be to fix a subset of these parameters (e.g., *θ*_2_ in our case) to a biologically realistic value, estimate the other parameters using CyGMM, and check the sensitivity of the results to changes in the fixed value as we did here.

## Acknowledgements

We would like to thank Ali Snedden and the high performance computing center (HPC) at the Nationwide Children’s Hospital for help with computation and John Whittman for his preliminary work on this project.

## Supplementary Information

This section is outlined as follows. First, we provide results from the estimation (and the PSO) when the analysis involves 10 proteins and 10 rate constants. Second, we offer compelling evidence that our nonlinear model is partially identifiable. Third, we give miscellaneous numerical details concerning the simulation of extrinsic noise, some of the hyper-parameters used in the PSO, and analysis of the real data.

### 5.1 Ten Proteins and Ten Rate Constants

As a proof of principle that CyGMM can handle a larger number of proteins, we generated 5 replicates of synthetic snapshot data (with 5,000 cells per replicate) from a linear ODE model with 10 proteins and 10 rate constants (1). Then, for each replicate, we used CyGMM to compute 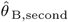, which incorporates means and variances with the optimal weights (Table 5).

**Table 5.** Estimates for 10-Protein Linear Model Estimated rate constants are shown for 5 independent replicates (denoted Rep) of synthetic snapshot data. The final row shows (in bold) the ground truth for comparison. The (*) indicates that multiple PSO swarms were needed to attain convergence. Each swarm takes about 5 minutes.

| Rep | $\theta_1$ | $\theta_2$ | $\theta_3$ | $\theta_4$ | $\theta_5$ | $\theta_6$ | $\theta_7$ | $\theta_8$ | $\theta_9$ | $\theta_{10}$ |
| --- | --- | --- | --- | --- | --- | --- | --- | --- | --- | --- |
| 1 | 0.531 | 0.051 | 0.108 | 0.925 | 0.056 | 0.058 | 0.195 | 0.762 | 0.360 | 0.223 |
| 2 | 0.500 | 0.028 | 0.065 | 0.880 | 0.039 | 0.038 | 0.156 | 0.746 | 0.310 | 0.253 |
| 3 | 0.500 | 0.123 | 0.176 | 0.900 | 0.132 | 0.136 | 0.211 | 0.768 | 0.360 | 0.222* |
| 4 | 0.504 | 0.046 | 0.095 | 0.935 | 0.075 | 0.072 | 0.176 | 0.741 | 0.334 | 0.217 |
| 5 | 0.508 | 0.055 | 0.100 | 0.899 | 0.060 | 0.073 | 0.143 | 0.706 | 0.330 | 0.225 |
|  | **0.516** | **0.060** | **0.103** | **0.897** | **0.055** | **0.072** | **0.177** | **0.747** | **0.343** | **0.232** |

To give some indication of how much PSO error there is from one swarm to the next, a single replicate was analyzed twice with two independent swarms. Shown below are the 10 estimated rate constants along with their corresponding costs:

0.531 0.051 0.108 0.925 0.056 0.058 0.195 0.762 0.360 0.223 **0.002**

0.529 0.083 0.144 0.922 0.092 0.085 0.202 0.771 0.367 0.222 **0.002**

This level of PSO error is commonly seen in about 80 - 90% of our 10-protein synthetic snapshot data sets. In contrast, shown below is an example of relatively high PSO error,

0.488 0.490 0.497 0.892 0.507 0.497 0.208 0.766 0.371 0.219 **0.022**

0.499 0.155 0.175 0.897 0.146 0.172 0.182 0.738 0.349 0.224 **0.006**

0.505 0.039 0.065 0.903 0.034 0.072 0.159 0.719 0.330 0.229 **0.001**

where the estimate with the minimum cost across 3 independent swarms is shown in the final row. This level of PSO error occurs in about 10-20% of all 10-protein snapshot data sets.

### 5.2 ODEs Used in the Nonlinear Model and for T cell Signaling Kinetics

#### 1. Nonlinear Model

The following ODEs describe time evolution of single cell protein abundances 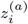 for the protein species *i* ≡ *Syk, V av*1, *Syk* − *V av*1, *SHP* 1, *pVav*1, *SHP* 1 − *pV av*1 in the model shown in Fig. 1D.

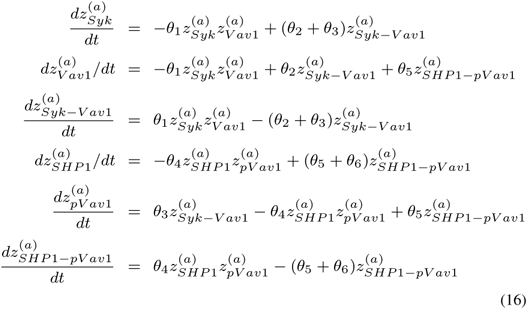

#### 2. T cell signaling kinetics

The following ODEs describe time evolution of single cell protein abundances 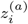 for the protein species *i* ≡ *pCD*3*ζ, pSLP* 76, *pErk, pS*6:

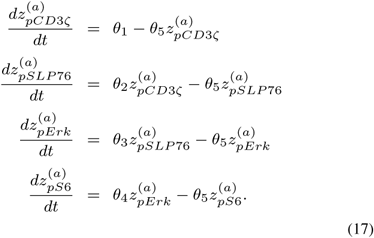

Pairwise contour plots of consecutive rate constants computed from protein abundances *pCD*3*ζ, pSLP* 76, *pErk*, and *pS*6 measured in T cells (Figure 4), provide multiple 2-dimensional perspectives of the actual 5-dimensional cost function. Furthermore, the contour plots are consistent with the hypothesis of a unimodal 5-dimensional cost function whose minimum value is near (1.541, 0.042, 0.311, 1.122, 0.384).

### 5.3 Partially Identifiable Models

Upon closer examination of our 6-protein nonlinear model, we found strong evidence that this model is partially identifiable (e.g. only 5 of the 6 rate constants can be estimated consistently). Table 6 shows the GMM estimate 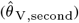 and its corresponding cost for different values of *θ*_2_. From a visual inspection, two things are immediately clear. First, the cost is an increasing function of *θ*_2_, which means that the estimate of *θ*_2_ is converging to zero. Second, *θ*_1_ depends linearly on *θ*_2_ (i.e. *θ*_1_ ≈ 0.09 + 0.10 ∗ *θ*_2_). Because the true value of *θ*_2_ is 0.10, the first point implies that *θ*_2_ cannot be estimated consistently. And the second point provides additional justification for fixing *θ*_2_ at 0.10, so that the estimation can focus on the remaining (presumably identifiable) rate constants. Note that, even when *θ*_2_ is incorrectly specified at 0.05 and 0.25, the estimates of the remaining rate constants are quite close to their true values. Furthermore, the value of the cost function changes very little as *θ*_2_ varies from 0 to 1.

**Table 6.** Evidence for a Partially Identifiable Nonlinear Model Estimated rate constants are shown for 13 different values of *θ*_2_. For each value, synthetic snapshot data were generated across 25,000 cells. Cost is shown in bold.

| $\theta_1$ | $\theta_2$ | $\theta_3$ | $\theta_4$ | $\theta_5$ | $\theta_6$ | Cost |
| --- | --- | --- | --- | --- | --- | --- |
| 0.091 | 0.010 | 0.945 | 0.170 | 0.051 | 0.180 | **0.00382** |
| 0.095 | 0.050 | 0.944 | 0.170 | 0.052 | 0.179 | **0.00391** |
| 0.100 | 0.100 | 0.945 | 0.170 | 0.051 | 0.180 | **0.00393** |
| 0.105 | 0.150 | 0.945 | 0.170 | 0.050 | 0.180 | **0.00410** |
| 0.115 | 0.250 | 0.946 | 0.170 | 0.050 | 0.180 | **0.00464** |
| 0.124 | 0.350 | 0.945 | 0.171 | 0.052 | 0.180 | **0.00525** |
| 0.134 | 0.450 | 0.945 | 0.170 | 0.051 | 0.180 | **0.00579** |
| 0.144 | 0.550 | 0.944 | 0.170 | 0.051 | 0.180 | **0.00645** |
| 0.153 | 0.650 | 0.944 | 0.170 | 0.051 | 0.179 | **0.00708** |
| 0.163 | 0.750 | 0.945 | 0.170 | 0.052 | 0.179 | **0.00769** |
| 0.173 | 0.850 | 0.945 | 0.170 | 0.051 | 0.180 | **0.00828** |
| 0.183 | 0.950 | 0.944 | 0.170 | 0.051 | 0.180 | **0.00885** |
| 0.186 | 0.990 | 0.944 | 0.169 | 0.050 | 0.179 | **0.00912** |

**Table 7.** Summary Statistics: Extrinsic Noise Summary statistic for the multivariate log-normal distribution are shown for the nonlinear ODE model with 6-parameters and 6-proteins. This is an example of a distribution that was used to simulate extrinsic noise in protein abundances.

| Statistic | Syk | Vav1 | Syk-Vav1 | pVav1 | SHP1 | SHP1-pVav1 |
| --- | --- | --- | --- | --- | --- | --- |
| Mean | 205.576 | 2059.205 | 5.310 | 5.411 | 205.535 | 5.309 |
| Median | 190.703 | 1997.600 | 5.271 | 5.396 | 190.630 | 5.268 |
| Q1 | 146.685 | 1693.801 | 4.855 | 5.120 | 146.924 | 4.852 |
| Q3 | 247.980 | 2356.792 | 5.723 | 5.684 | 247.431 | 5.722 |

**Table 8.**
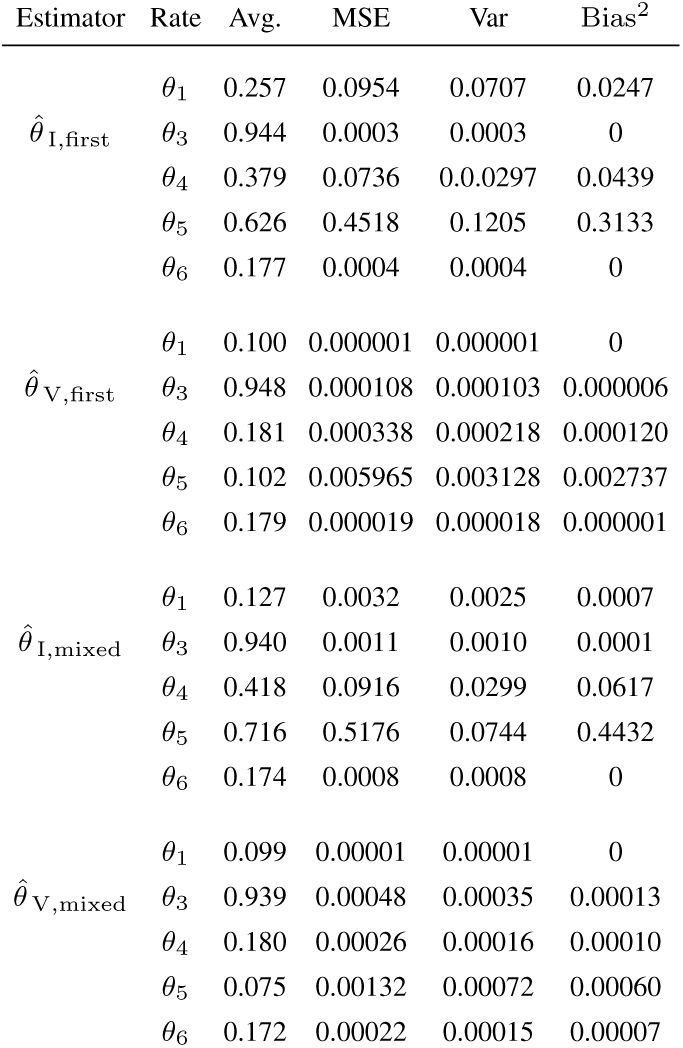
MSE (Nonlinear Model with 6 Proteins) The average across 100 replicates, MSE, variance, and squared bias are shown for four different GMM estimators. Two estimators use only the difference in mean protein abundance, and two estimators use differences in means, variances, and covariances. For each replicate, *θ*_2_ was fixed at 0.10, and the remaining rate constants were estimated with our GMM approach.

### 5.4 Miscellaneous Numerical Details

We generate synthetic snapshot data with extrinsic noise using a two step procedure. For each cell, we first sample values from a multivariate normal distribution. This allows (if desired) the abundances of some (possibly all) proteins to be correlated. Then, in the second step, each element of that random vector is exponentiated, so that the resulting abundances are marginally log-normal. Table (7) shows the mean, median, and inter-quartile range (IQR) for the nonlinear model, which is loosely based on the biochemical signalling biology reported in (Das (2010)).

From Figure 5, we can see how the contours are more concentrated when means and variances are used in the cost function (right), compared to contours where the cost function just uses differences in means (left). This is consistent with Table (3) which shows (among other things) that estimation is improved by including higher order moments.

**Figure 5.**
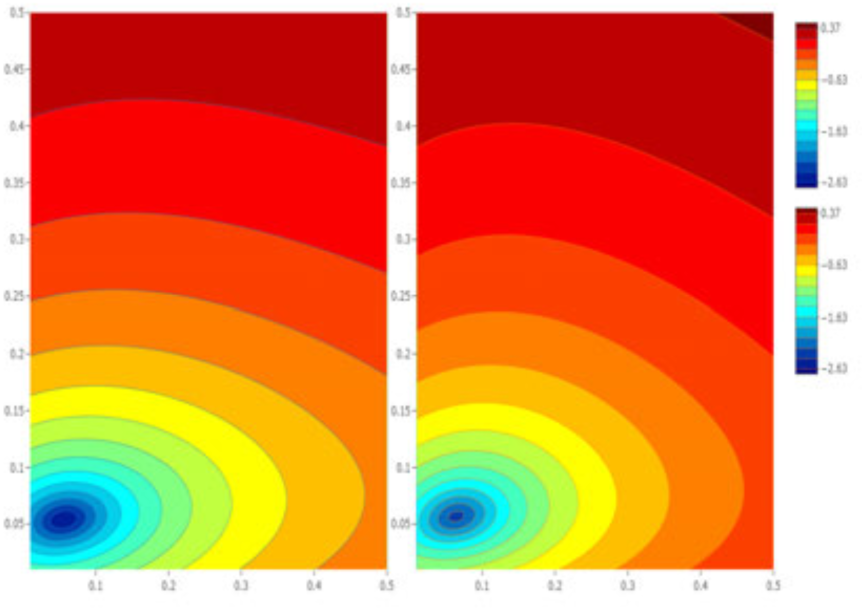
Two pairwise contour plots showing increased precision for parameter estimation when higher order moments are included. The high dimensional cost function in the 6-parameter 6-protein linear ODE model is shown with differences in first moments considered (Panel A), and differences in first and second moments considered (Panel B). For each contour plot, optimal weights were used; the x-axis shows *θ*_2_ and the y-axis shows *θ*_5_. The remaining rate constants were fixed at their true values.

